# A Comparative Analysis of Imaging-Based Spatial Transcriptomics Platforms

**DOI:** 10.1101/2023.12.13.571385

**Authors:** David P. Cook, Kirk B. Jensen, Kellie Wise, Michael J. Roach, Felipe Segato Dezem, Natalie K. Ryan, Michel Zamojski, Ioannis S. Vlachos, Simon R. V. Knott, Lisa M. Butler, Jeffrey L. Wrana, Nicholas E. Banovich, Jasmine T. Plummer, Luciano G. Martelotto

## Abstract

Spatial transcriptomics is a rapidly evolving field, overwhelmed by a multitude of technologies. This study aims to offer a comparative analysis of datasets generated from leading *in situ* imaging platforms. We have generated spatial transcriptomics data from serial sections of prostate adenocarcinoma using the 10x Genomics Xenium and NanoString CosMx SMI platforms. Additionally, orthogonal single-nucleus RNA sequencing (snRNA-seq) was performed on the same FFPE tissue to establish a reference for the tumor’s transcriptional profiles. We assessed various technical aspects, such as reproducibility, sensitivity, dynamic range, cell segmentation, cell type annotation, and congruence with single-cell profiling. The practicality of assessing cellular organization and biomarker localization was evaluated. Although fewer genes are measured (CosMx: 960, Xenium: 377, with an overlap of 125), Xenium consistently demonstrates higher sensitivity, a broader dynamic range, and better alignment with single-cell reference profiles. Conversely, CosMx’s out-of-the-box segmentation outperformed Xenium’s, resulting in noticeable transcript misassignment in Xenium within certain tissue areas. However, the impact of this on the cells’ transcriptional profile was minimal. Together, this comprehensive comparison of two leading commercial platforms for spatial transcriptomics provides essential metrics for assessing their performance, offering invaluable insights for future research and technological advancements in this dynamic field.

## Introduction

Single cell/nucleus RNA sequencing (sc/nRNA-seq) has facilitated numerous scientific breakthroughs, enabling researchers to identify rare cells^1^, delineate dynamic cell states^2^, and decode cellular communication networks within complex biological systems^3^. Its advantage over bulk RNA sequencing (RNA-seq) lies in the preservation of individual cell transcriptomes, avoiding the homogenization of transcripts from all cells in a sample. However, challenges persist with snRNA-seq for various tissue types, and the loss of spatial information of individual cells hinders the full elucidation of complex systems and disease states.

Spatial transcriptomics (ST) technologies overcome the limitations of snRNA-seq by enabling large-scale *in situ* profiling of RNA transcript distributions within tissue sections. Imaging-based ST (iST) methods involve the preparation of tissue cross-sections for repeated hybridization with fluorescently labeled probes that bind to specific mRNAs; this is followed by microscopy imaging to capture the spatial distribution of transcripts^4^. The integration of individual cell transcriptomes with their spatial context promises to revolutionize our understanding of biological processes. Furthermore, iST platforms capable of analyzing formalin-fixed, paraffin-embedded (FFPE) samples will open access to a wealth of historical clinical pathology specimens preserved in archival tissue banks.

iST methods, however, are not without their own challenges. Probes and fluorescence imaging must be sufficiently specific to avoid false positives from off-target binding or autofluorescence, and must reliably bind to and signal from their intended transcripts to reduce false negatives. FFPE samples present particular difficulties due to RNA degradation^5^. Achieving high sensitivity and specificity is crucial for accurate cell typing, and these challenges are particularly pronounced when dealing with FFPE tissue microarrays (TMAs), as was recently underscored^6^.

In this study, we undertake an in-depth comparative analysis of two leading *in situ*, imaging-based spatial technologies: the Xenium platform from 10x Genomics ^7^ and the CosMx Spatial Molecular Imager from NanoString Technologies^8^. We base our comparison on data gathered from serial sections of a prostate cancer FFPE sample. Comparisons such as this are crucial, as these technologies represent the forefront of digital pathology tools and will become instrumental in advancing our understanding of complex biological systems. To establish a robust benchmark for transcript quantifications, we also generated single-nucleus RNA sequencing data from the same tissue block using an established method for processing formalin-fixed paraffin-embedded (FFPE) samples^9^—hereafter *snPATHO-seq*. Our findings indicate that despite their obvious similarities, the two platforms differ significantly across all metrics. CosMx presents increased assay plexity and improved cell segmentation capabilities, but it is hindered by lower sensitivity compared to Xenium. The latter is manifested by many expressed genes not detected above background levels, sparse detection of cell type markers, and limited consistency with the orthogonal snRNAseq data. This ultimately affects downstream analyses, including cell type annotation and biological investigations that depend on the reliable detection of a limited number of genes, such as biomarker discovery, cell state dynamics, and cell communication inference. Benchmarking iST platforms enhances our methodological arsenal and is an essential step in comprehending the optimal application of these technologies in cancer research and broader scientific inquiries^6^.

## Results

### Processing matched prostate cancer samples with Xenium, CosMx, and snPATHO-seq

To facilitate a direct comparison of data from the Xenium and CosMx platforms, we designed an experiment to assess serial sections of a primary prostate adenocarcinoma (**Figure 1A**). Two consecutive sections were processed using each platform’s commercial iterations, utilizing pre-designed probe sets for general tissue profiling (a human 1,000-plex gene panel for CosMx; a human 377-plex multi-tissue and cancer panel for Xenium). A single section from in-between was stained with Hematoxylin and Eosin (H&E) for histological analysis. Furthermore, to establish a matched reference for the transcriptional profiles of cell types within the tumor, four adjacent sections underwent fixed RNA profiling from single nuclei (snPATHO-seq) ^9^.

**Figure 1.**
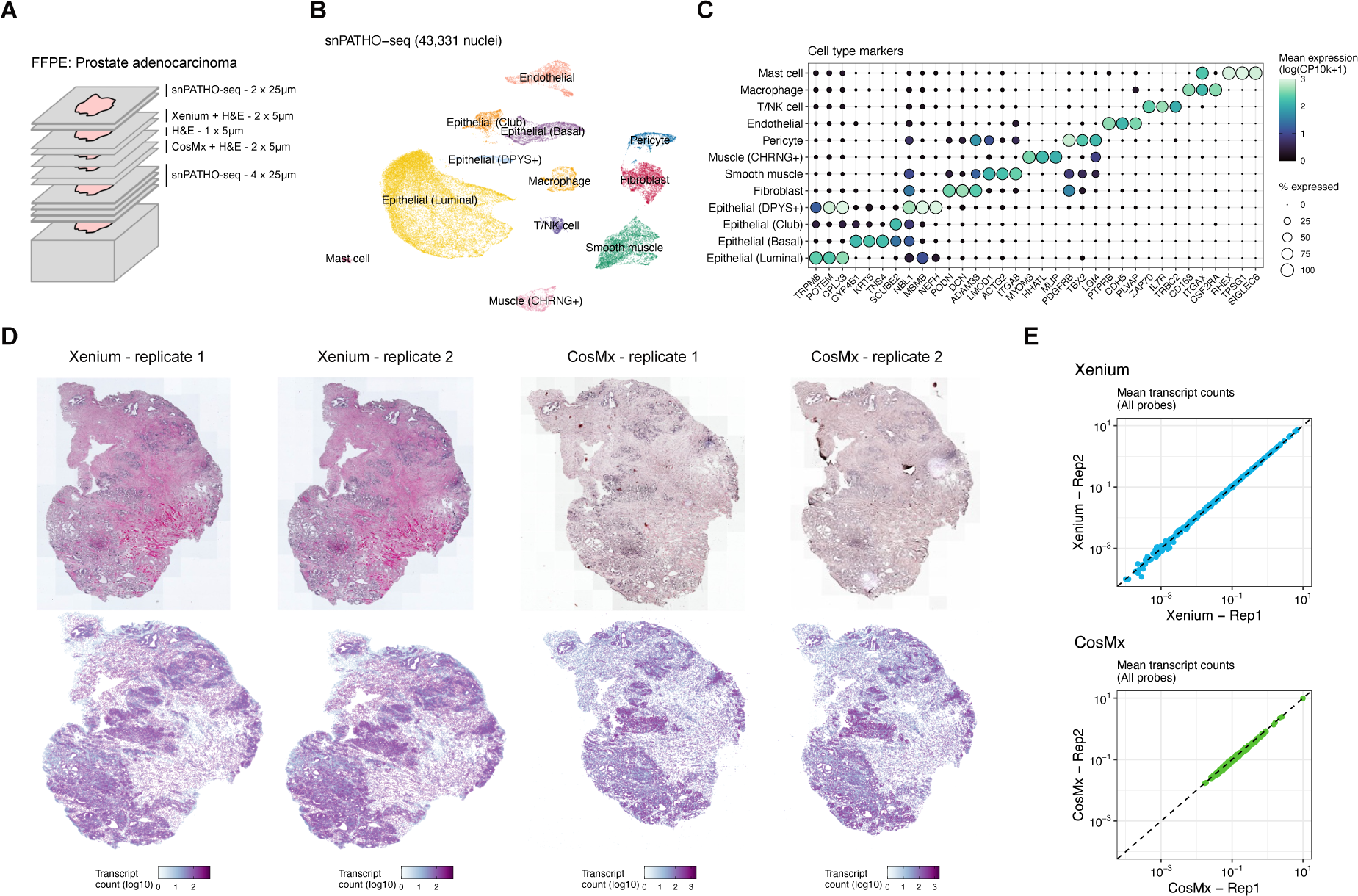
**A.** Schematic of the Experimental Design: Serial sections of FFPE prostate adenocarcinoma were simultaneously processed for spatial transcriptomics using Nanostring CosMx SMI and 10x Genomics Xenium, single-nucleus RNA sequencing (snPATHO-seq), and histological analysis. **B.** Annotated UMAP Embedding of snPATHO-seq Data: Cell types were annotated utilizing reference scRNA-seq data of human prostate from Tuong et al. ^10^, along with canonical markers for each cell type. **C.** Expression and Detection Frequency of Top 3 Markers: The top three markers (p<0.05, ranked by logFC) for each cell type are shown. Log(CP10k+1) represents the log-transformed transcript counts per 10,000 transcripts. **D.** Top: H&E Stain of Each Section Following Xenium or CosMx Imaging: Post-imaging, CosMx tissues undergo additional processing for immunofluorescence, which affects the stain quality. Bottom: Spatial Transcriptomics Data: Each cell is depicted as a point, positioned at its centroid. Cells are color-coded by total transcript count. **E.** Correlation of Mean Transcript Counts for Each Probe in the Assay Across Replicates on the Same Platform: The Pearson correlation coefficient for both platforms >0.99.

snPATHO-seq recovered 43,331 high-quality nuclei from the tissue (**Figure 1B**). Using an annotated scRNA-seq dataset from human prostate as a reference^10^, all major cell types were present in the sample (**Figure 1B,C**). This includes a prominent luminal epithelial population representing the tumor’s malignant component^11^, as well as various other epithelial lineages such as club, basal, and a DPYS+ luminal population. All major non-epithelial components, including lymphocytes, endothelial, and muscle cells were also identified (**Figure 1B,C**). Histology of the sections processed on Xenium and CosMx were similar, and the cellular arrangement captured in the spatial transcriptomics data reproduced the tissue structure (**Figure 1D**). Additionally, the average transcript quantifications between replicate sections were highly reproducible on both platforms (**Figure 1E**).

### Xenium achieves higher sensitivity and dynamic range than CosMx

Both CosMx and Xenium platforms detected approximately 100,000 cells (ranging from 96,139 to 102,508) in each of the four tissue sections analyzed (**Figure 2A**). The CosMx probe set targets over twice as many genes compared to Xenium’s (960 vs. 377, respectively), resulting in a proportionally higher number of genes detected per cell in the two CosMx sections. However, when accounting for differences in assay plexity–the number of gene targets in a probe set–we found that the median number of transcripts detected per gene was two-fold higher in the Xenium data (**Figure 2A**). Importantly, when limited to the 125 genes targeted by both probe sets, Xenium recovered approximately three times more transcript counts per cell than CosMx (**Figure 2A,B**).

**Figure 2.**
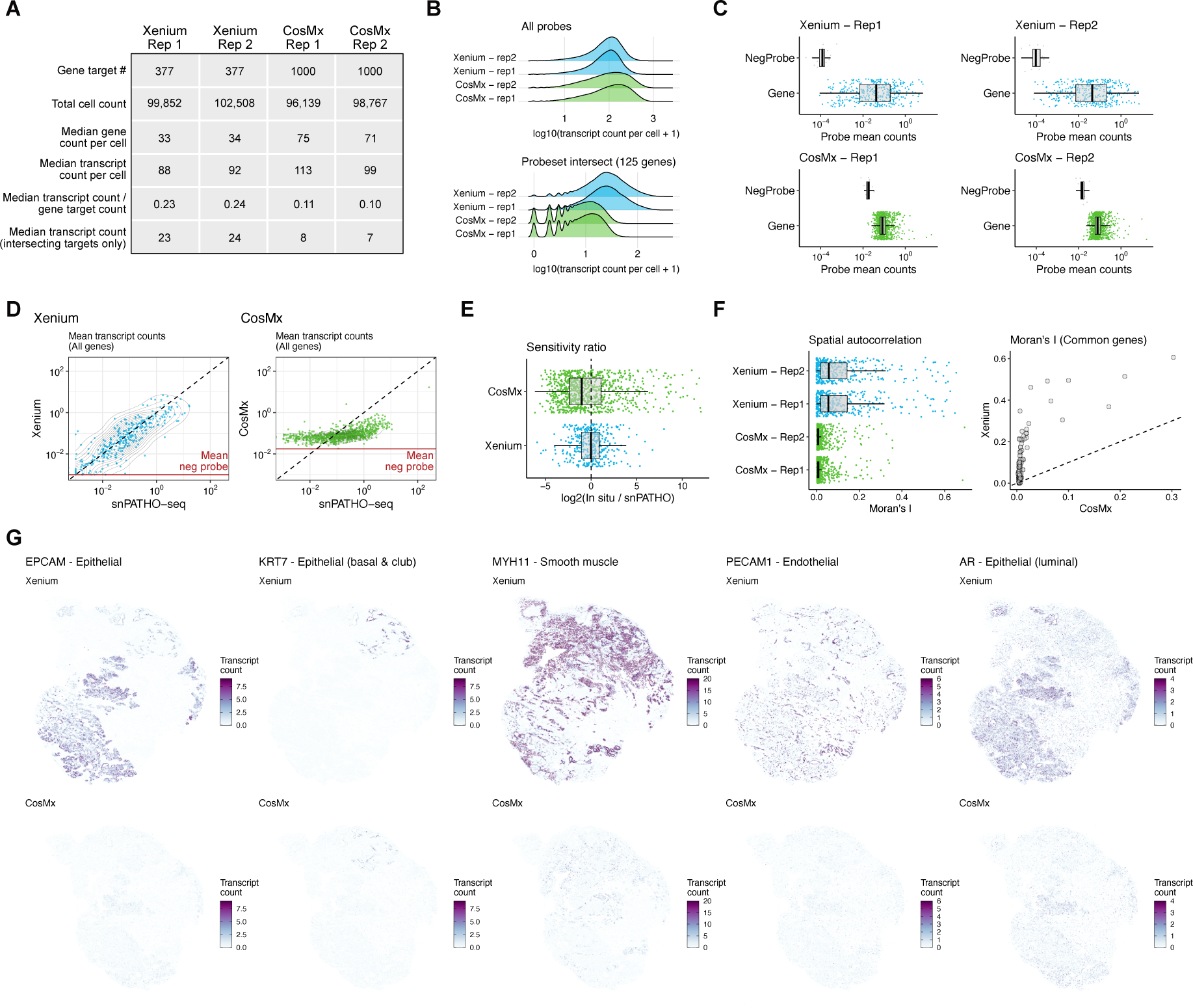
**A.** Summary of various quality control metrics for each sample. **B.** Distribution density of total transcripts detected per cell across all probes in each assay (top) and focusing solely on the 125 probes in each assay targeting common genes (bottom). **C.** Mean transcript counts for all gene-targeting probes (“Gene”) and negative control probes (“NegProbe”) in each sample, with boxes and whiskers representing the 25th-75th percentiles and 1.5x the interquartile range (IQR), respectively. **D.** Correlation between average transcript counts in Xenium (left) and CosMx (right) samples with snPATHO-seq data, displaying all gene-targeted probes. The dashed line indicates a slope of 1, while the solid red line represents the average abundance of the negative control probes for each platform. **E.** Log2 ratio of average transcript counts in the spatial transcriptomics data compared to the snPATHO-seq data. **F.** Left: Spatial autocorrelation (Moran’s I) of all genes targeted in each assay. Right: Comparison of Moran’s I values for genes targeted by both platforms, consistently higher in Xenium data than in CosMx. **G.** Per-cell transcript counts for several cell type markers in the first replicate of both Xenium (top row) and CosMx (bottom row), with the colormap range manually set and values exceeding the limits colored as the maximal value.

To better understand the sensitivity and dynamic range of the two platforms, we evaluated the average transcript counts for gene and non-targeting control probes in each data set. The 20 control probes in the Xenium platform were consistently detected at lower rates than the 11 included with CosMx, though both were detected at quite low rates (mean <0.01 transcripts per cell) (**Figure 2C**). Gene-targeting probes for both platforms were consistently detected above the levels of control probes, though the median transcript abundance for Xenium was 372-fold over the median control probe abundance, but only 4.7-fold for CosMx. The dynamic range of Xenium quantifications was also higher, with a 4.3-fold higher interquartile range (Xenium: 6.9e-3 to 2.2e-1; CosMx: 0.6 to 0.1). We then compared transcript abundances from the two platforms to the matched snPATHO-seq. Although these counts were averaged across the entire population, we observed that the Xenium data correlated well with the snPATHO-seq data across its entire range of detection (**Figure 2D,E**). In contrast, the CosMx data exhibited a noticeable inflation of lowly expressed genes, contributing to its compressed dynamic range. As a result, genes varying three orders of magnitude in the snPATHO-seq data had similar detection in the CosMx data. This is compounded by a reduced sensitivity for highly expressed genes, which were recovered at lower levels than observed in the snPATHO-seq (**Figure 2D,E**).

Cellular organization throughout tumors is notably heterogeneous^12^. Prostate adenocarcinomas are characterized by glandular outgrowths of malignant cells that invade the smooth muscle-rich stroma of the prostate. These glandular structures remain physically distinct from stromal components (**Figure 1D**). Given this inherent structure, many cell type-specific transcripts are expected to be concentrated in distinct regions throughout the tumor. Using the spatial autocorrelation metric Moran’s I, we found that spatial patterning of genes throughout the tissue was more prominent in the Xenium sample (**Figure 2F**). Consistent with this, we found that the abundance and specificity of cell type-specific transcripts to various histological structures was notably higher in the Xenium data (**Figure 2G**).

### Poor sensitivity impacts cell type annotation

To evaluate the ability to discern cellular organization we annotated cell types in the prostate cancer data using the snPATHO-seq data as a reference for automated annotation. For all samples, histology was broadly resolved and showed general consistency across both platforms (**Figure 3A**). However, in the CosMx samples, we observed a higher incidence of cells with epithelial annotations throughout the tumor stroma, despite the lack of transcripts associated with epithelial markers (**Figure 3B**).

**Figure 3.**
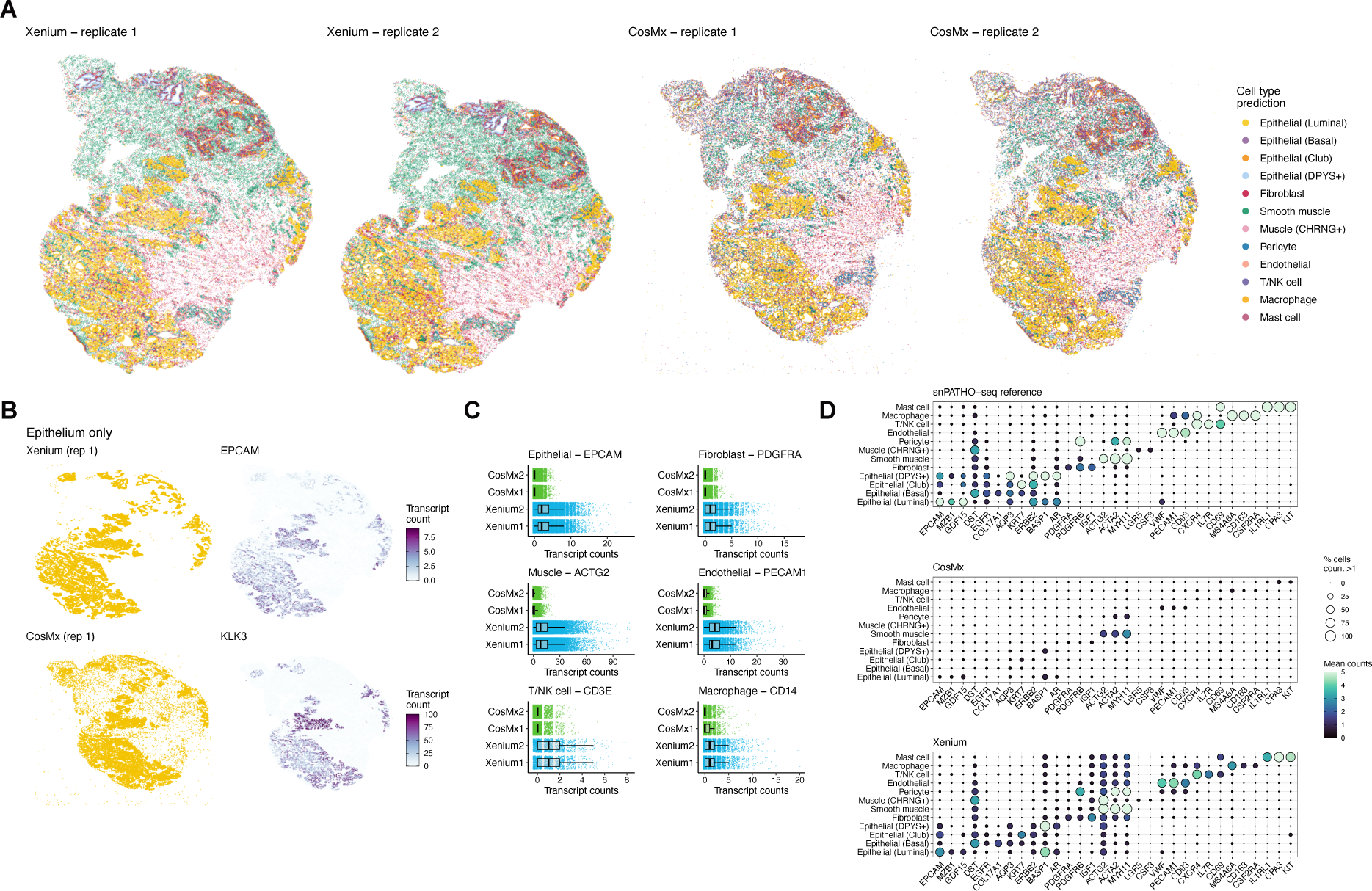
**A.** Annotations of cell types in spatial transcriptomics data. **B.** Left: Distribution of cells identified as epithelial. Unlike in Xenium data, CosMx samples show a higher prevalence of epithelial annotations in stromal areas. Right: Patterns of expression for epithelial markers (*EPCAM* in Xenium; *KLK3* in CosMx) across the tissue. Notably, despite the annotations in CosMx, there is no *KLK3* expression in the tumor stroma. **C.** Detection frequencies of canonical cell type markers, limited to cells with corresponding annotations. **D.** Analysis of cell type marker detection and abundance in snPATHO-seq, CosMx, and Xenium data. These markers were selected based on their differential expression in snPATHO-seq data (Wilcoxon rank-sum test, p < 0.05, top 3 ranked by log fold change). The color map’s range was set manually, and values beyond the limits were not shown.

Misidentification may be linked to the lower signal-to-noise ratio exhibited by CosMx, affecting the detection of stromal cell transcripts. To investigate this, we analyzed the abundance of marker transcripts for various annotated cell types and observed that many canonical markers were detected at lower abundances in the CosMx data (**Figure 3C**). We further conducted a systematic evaluation using snPATHO-seq data to initially define cell type markers for this specific tumor. In this analysis, cells in the Xenium dataset consistently showed a higher detection rate and dynamic range for all markers compared to those in the CosMx samples (**Figure 3D**). However, it is important to note that Xenium exhibited a higher incidence of unrelated marker detection, especially among stromal cell types, potentially indicating transcript contamination from imperfect segmentation.

### Xenium’s higher sensitivity and dynamic range is consistent in independent samples

To determine whether the differences observed between platforms are consistent across independent samples, we next compared available Xenium and CosMx data from several tissue types, including human colon, lung cancer, and breast cancer. For each tissue type, we also compared transcript quantifications to reference single-cell RNA sequencing (scRNA-seq) data from unmatched samples. It is important to note that only the breast cancer samples were derived from matched FFPE tissue, and all samples were processed at different times in different laboratories.

In each tissue type, we observed common patterns similar to those in the prostate cancer data, albeit to varying extents. Gross tissue structure could be resolved after automated annotation using a scRNA-seq reference (**Figure 4A**). Negative control probes were consistently detected at higher levels in the CosMx and many gene-targeting probes were only modestly detected above this threshold (**Figure 4B**). The Xenium transcript quantifications span a wider dynamic range and higher spatial autocorrelation (**Figure 4B,C**). Correlation of CosMx data with public scRNA-seq data was better than we observed in the prostate cancer data, though lowly expressed genes were similarly inflated and there is a tendency towards lower detection of highly expressed genes (**Figure 4D**). Related, canonical marker expression is often higher in Xenium data (**Figure 4E**). The consistent observation of these patterns suggests that they may be inherent properties of the platforms rather than consequences of tissue processing.

**Figure 4.**
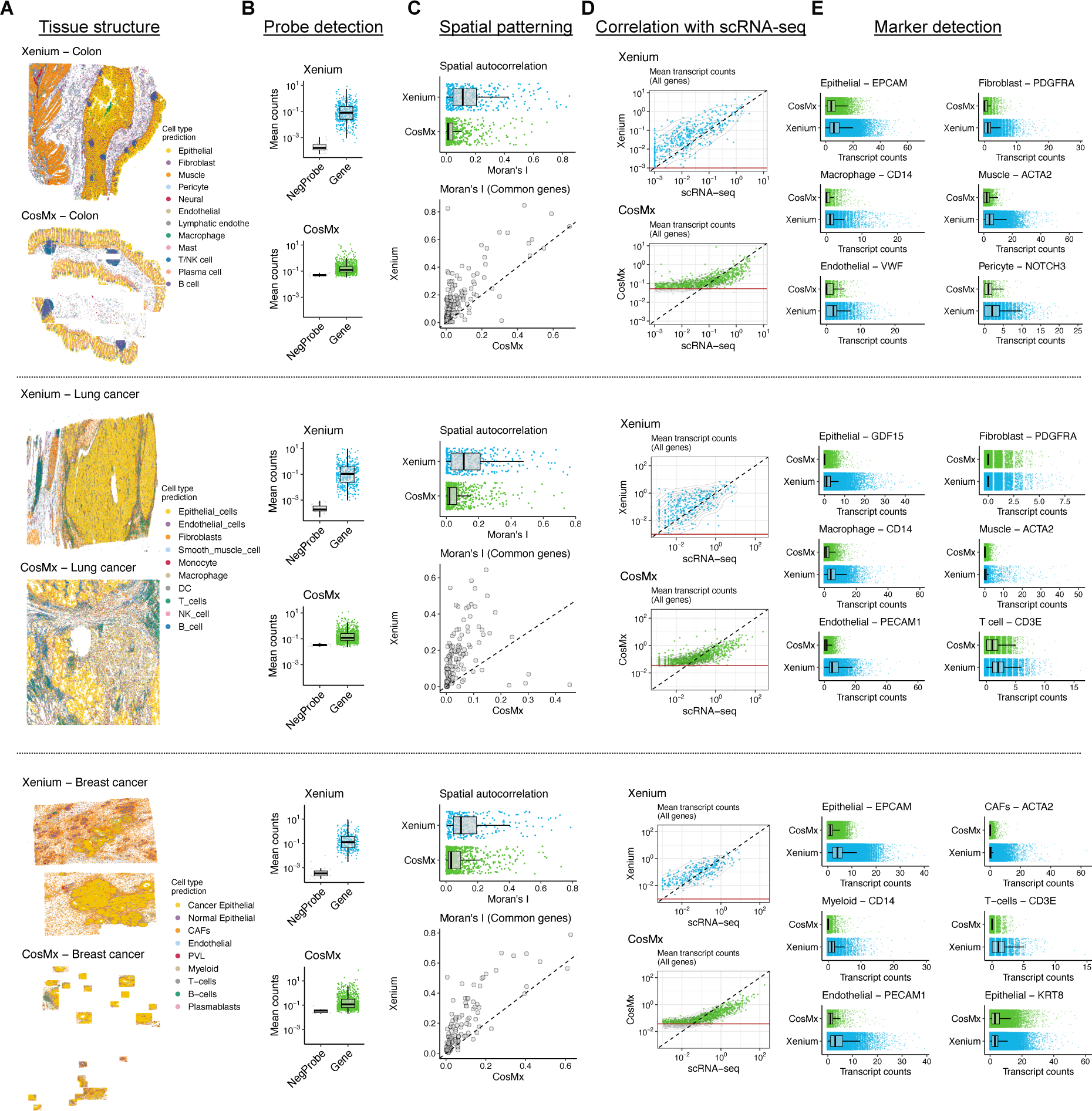
**A.** Tissue structure of colon (top; Xenium: public data from 10x Genomics, CosMx: Garrido-Trigo et al.^23^), lung cancer (middle, Xenium: public data from 10x Genomics, CosMx: public data from NanoString Technologies), and breast cancer (bottom; Xenium and CosMx provided the Suzuki lab, University of Tokyo). Cell annotations are based on automated annotation from scRNA-seq reference datasets. **B.** Average detection of gene-targeting and control probe sets. **C.** Spatial autocorrelation (Moran’s I) of all probes in each data set (top panels). A comparison of Moran’s I values for common gene targets is shown in each bottom panel. The dashed line reflects a slope of 1. **D.** Correlation of spatial transcriptomics data from each platform with a scRNA-seq reference (Breast cancer, Wu et al. ^24^; Lung cancer, Kim et al. ^25^; Colon, custom integration of public data, available with data from this manuscript). Given that reference data is from a different source and cellular composition may differ, values only from annotated endothelial cells are shown as an example. **E.** Transcript abundance for canonical markers in cells with the specified annotation are colored as the maximal value.

### Segmentation and assay sensitivity impact the ability to resolve cellular organization

In our comparison of the spatial transcriptomics and snPATHO-seq data, we observed a higher frequency of unrelated marker detection, notably among stromal cell types in the Xenium data. For instance, when compared to snPATHO-seq data, smooth muscle markers such as *ACTG2* and *ACTA2* were consistently identified at higher rates in other stromal cell types (e.g., immune cells) (**Figure 4D**). As the transcriptional profiles are dependent on the cell segmentation, we further explored how segmentation might contribute to potential misassignment of transcripts. We focused on a tissue region containing a network of blood vessels (**Figure 5A**). Histologically, these vessels are lined by a thin layer of endothelial cells enveloped by pericytes. Each vessel is surrounded by a layer of smooth muscle and located within a network of fibroblasts and a protein-rich ECM.

**Figure 5.**
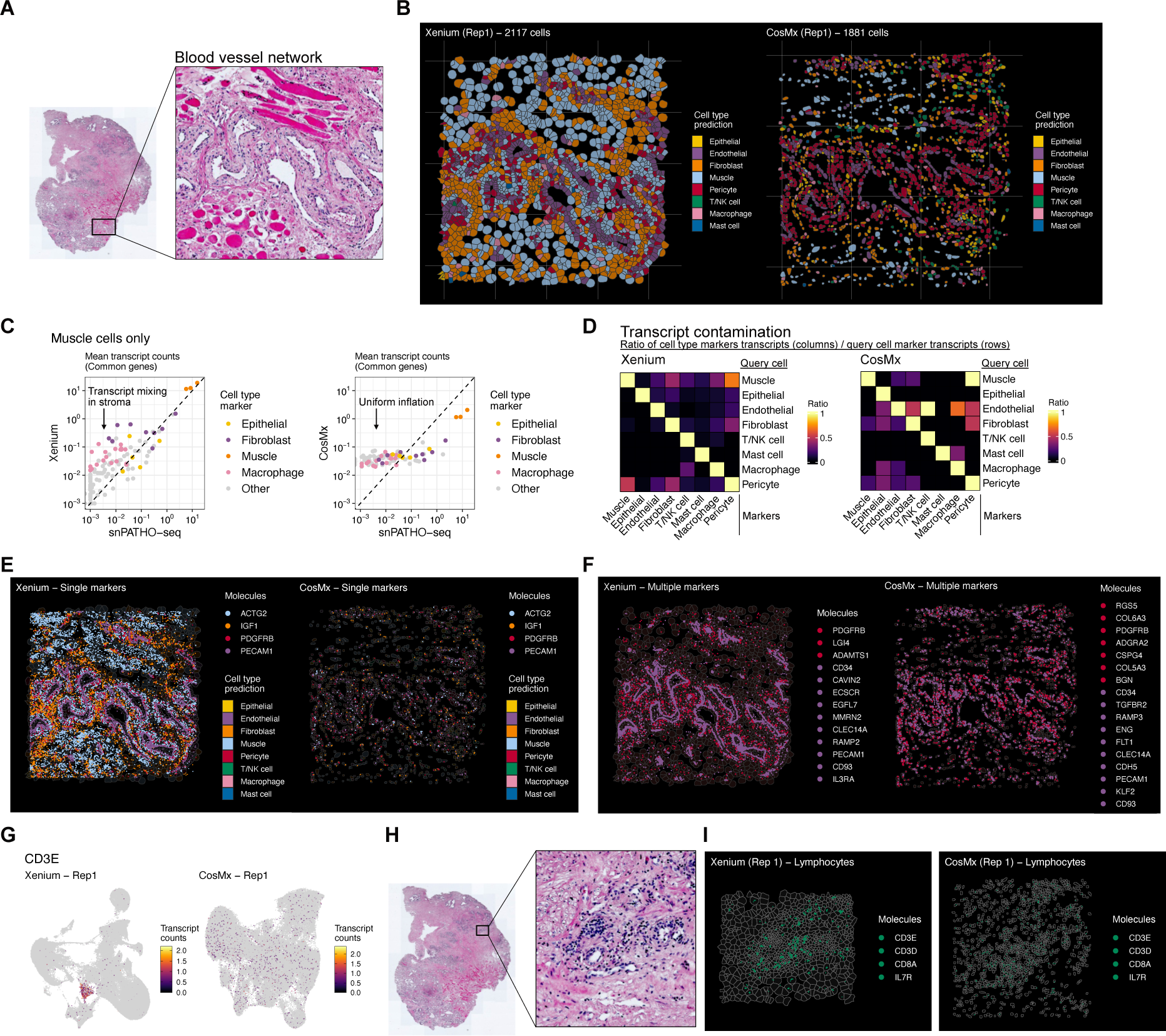
**A.** H&E stain of a tissue region with prominent blood vessels and structured supportive cell types (smooth muscle, skeletal muscle, pericytes). **B.** Cell segmentation of Xenium and CosMx data in an overlapping region to (A). **C.** Average transcript counts of muscle cells in snPATHO-seq, Xenium, and CosMx data. Markers of select cell types are indicated. Values above the dashed line reflect *in situ* quantifications higher than those in the snPATHO-seq. **D.** Heatmaps showing the median ratio of transcripts for unrelated cell types to those of the cells’ annotated identity. The plot shows the relative abundance of markers (columns) in various cell types (rows). **E.** Same region as (B) showing the abundance and localization of individual markers for muscle (*ACTG2*), fibroblasts (*IGF1*), pericytes (*PDGFRB*), and endothelial cells (*PECAM1*). Cells with >5 transcripts: Xenium, 368; CosMx, 7 **F.** Same as (E), except showing all Re tv endothelial and pericyte markers (p<0.05, logFC>1, pct.out < 20) targeted in either probe set. **G.** UMAP embedding of Xenium and CosMx data. Cells are coloured by the number of *CD3E* transcripts. **H.** H&E stain of a tissue region with abundant lymphocytes. **I**. Overlapping region with (H) showing detection of *CD3D/E*, *GZMA*, and *IL7R*. Cells with >5 transcripts: Xenium, 62; CosMx, 1.

When analyzing this region, we observed that the default segmentation algorithm applied to Xenium data inflates cell boundaries, resulting in artificially enlarged cell sizes, especially in sparsely populated areas (**Figure 5B**). This algorithm is based on expanding nuclear boundaries (determined by DAPI stain), which differs from CosMx’s approach, which utilizes cell membrane staining. Based on cell size and arrangement, CosMx’s default segmentation more faithfully replicates cell morphology. Nevertheless, there are challenges in accurately defining the boundaries of cells with complex, narrow, or elongated shapes using either platform.

To evaluate how inaccurate segmentation might produce mixed transcriptional profiles, we investigated the stroma, known for its heterogeneous composition. Given the diffuse distribution of muscle cells throughout the tumor’s stroma, we explored the potential misassignment of transcripts. In the Xenium data, macrophage and several fibroblast markers were more abundant than expected in annotated muscle cells based on snPATHO-seq data, whereas various epithelial markers were not (**Figure 5C**). To further assess the prevalence of unrelated markers in each cell type, we analyzed the relative abundance of off-target marker transcripts. Specifically, we first identified a list of markers for each cell type (p<0.05, top 10 genes ranked by logFC). We then calculated, for all cells of a given type, the ratio of transcripts from each marker set relative to the markers defining the cell’s identity. In the Xenium data, we observed markers for endothelial, fibroblast, and muscle cells in pericytes (**Figure 5D**), consistent with transcript contamination from neighboring cells. Markers for various stromal cell types were also detected in muscle cells, likely due to expanded cell boundaries. However, in the CosMx data, marker detection was generally less specific, which could be attributed to lower abundance of on-target markers and higher noise levels, rather than segmentation errors (**Figure 5D**).

The precise localization of cell type-specific transcripts can significantly enhance our understanding of cellular organization and is not influenced by segmentation challenges. To illustrate this, we examined markers of endothelial cells (*PECAM1*), pericytes (*PDGFRB*), fibroblasts (*IGF1*), and muscle (*ACTG2*). We found that tissue organization, consistent with histological findings, was clearly delineated in the Xenium data (**Figure 5E**). For example, despite the endothelial segmentation expanding into the vessel lumen, a thin layer of PECAM1 transcripts accurately highlighted the lumen structure. CosMx data, however, struggled to define spatial organization effectively due to the low detection rates of each marker (**Figure 5E**). When assessing multiple cell markers, however, the ability to discern tissue structure in the CosMx data was improved (**Figure 5F**).

Finally, we examined the impact of segmentation and assay sensitivity on the detection of rare cell types. Clustering the spatial transcriptomics data, we identified a population in the Xenium data that distinctly corresponded to CD8 T cells (CD3+, CD8A+, GZMA+) but was absent from the CosMx data (**Figure 5G**). The cell type annotations from the Xenium data revealed several regions in the tumor characterized by an increased abundance of T cells. In these areas, the Xenium data showed punctate regions with concentrated lymphocyte markers (**Figure 5H, I**). In contrast, while these markers were detected in the CosMx data, they were more uniformly distributed and not detected at levels sufficient to as confidently localize T cells (**Figure 5I**). The small size of lymphocytes and their low RNA content may make their detection particularly vulnerable to both segmentation and sensitivity issues. This underscores that the reduced sensitivity of CosMx may lead to a systematic limitation in its ability to detect specific cell types.

## Discussion

Our comparative analysis of Xenium and CosMx spatial transcriptomics platforms, augmented by snPATHO-seq data, offers valuable insights into the evolving field of spatial transcriptomics, transcending the specific context of cancer research and providing broader implications for various biological and medical research areas.

A key observation from this analysis has been the importance of assay sensitivity. Simply, many analyses depend on the robust detection of relevant transcripts. Although CosMx data has consistently higher noise that masks lowly expressed transcripts, the challenges that emerge in accurate cell typing and data interpretation tend to come from the weak detection of highly expressed cell type markers. Assuming sensitivity remains similar, increasing probe set plexity may improve the performance of these analyses, as they can be informed by a larger number of relevant genes. However, it is critical to underscore that many biological questions will depend on the accurate detection of a small number of genes: tumor classification based on the presence of a single biomarker, distinguishing between subtly polarized cell states, the spatial mapping of cells expressing immunosuppressive factors, and more.

The promise of simultaneously measuring thousands of genes is certainly appealing, and it is often argued that this is required to capture more biological processes. However, it is unclear if this can be accomplished without significantly affecting data quality. Optical resolution is finite, and the inclusion of more targets may reduce the ability to confidently detect individual transcripts. Increasing the number of probes may also increase the number of off-target binding events, elevating noise in the data. As demonstrated by the practice of “highly variable gene selection” in the analysis in scRNA-seq data, many biological processes involve variable expression of a limited number of genes (hundreds to low thousands). While large probe sets are a convenient off-the-shelf option, custom low-plex probe sets can be tailored to adequately capture relevant genes. And in most cases, accurate cell typing can be accomplished with a limited probe set. For example, although the probe set for the Xenium samples was a general tissue profiling panel of 377 genes, all cell types of the tissue could be identified, including four distinct epithelial subtypes.

Cell segmentation is a common feature compared between platforms. However, it is noteworthy that this is the one technical component of the data that is not inherent to the platform itself. Independent segmentation algorithms can be applied to data from all iST platforms, and as new tools are developed, this may become common practice. Neither platform currently employs algorithms that consider the localization and identity of the detected transcripts themselves, which has been proposed recently to improve segmentation^13,14^. The inclusion of protein detection to inform segmentation of the CosMx data is clearly beneficial, resulting in a more faithful representation of the cells’ morphology than Xenium. Though it is unclear if tissue processing for immunofluorescence prior to probe hybridization impacts transcript detection. Although Xenium’s default segmentation results in artificially inflated cell sizes, we found that this only results in marginal misassignment of transcripts from adjacent cells. This is likely because the most extreme cell sizes are from cells in sparsely populated regions where the nuclear expansion can proceed without encountering another cell boundary.

In summary, our study emphasizes that in spatial transcriptomics, the choice of platform should be guided by the specific research goals, with a particular focus on the sensitivity and dynamic range. These factors are critical for the accurate detection of gene expression patterns and cell types, which in turn are essential for drawing meaningful biological conclusions. As spatial transcriptomics continues to evolve, these insights will be instrumental in guiding both the development of new technologies and the design of experiments across a wide range of biological and medical research fields.

### Caveats of this study

The study presents several caveats that are important for understanding its context and implications:

1. Sample Size: The study, despite its limited sample size, marks an important first step in establishing benchmarking methods in spatial transcriptomics. It serves as foundational work for future studies, including those involving cross-institution comparisons, and will aid in better assessment and interpretation of emerging data.
2. Matching Cell Segmentations from H&E: A technical challenge encountered was aligning cell segmentations from H&E stained sections for accurate transcript assignment. Addressing this hurdle is crucial for enhancing the accuracy of spatial transcriptomics data^15,16^.
3. Comparison with Other Platforms: The scope of the study was limited to two leading specific platforms and did not include a range of other in situ imaging platforms. As more comprehensive data becomes available, the comparative methods used here can be extended to offer broader insights into the capabilities of various platforms.
4. Tissue Type-Specific Effects: The impact of tissue types on spatial transcriptomics results is a significant factor. Other studies, including recent research from the Broad Institute ^6^ and SciLab ^17^ have highlighted the importance of tissue-specific considerations in interpreting results. Computational tools are quickly evolving to address this ^18^.
5. Biological Significance of Transcripts in Undefined Cells: An empiric but intriguing observation is the presence of transcripts in undefined cell regions, referred by our team to as the “nebula,” across different platforms. The biological relevance of these transcripts remains uncertain, yet it’s crucial to consider their potential impact on the overall interpretation of the data.
6. Protein Acquisition and Multiomic Approaches: While protein measurements or combinations with other ‘omics (e.g. accessibility of chromatin) was beyond the scope, we acknowledge that future research integrating these elements is likely to yield deeper insights into cell annotation and contribute to the development of new benchmarking methods. This approach is expected to significantly deepen our understanding of cellular mechanisms and interactions.
7. Segmentation-free tools: Image-based cell segmentation in spatial transcriptomics faces challenges like misassigned RNA transcripts due to 3D to 2D tissue conversion,segmentation inaccuracies in non-spherical and special cell types, variability in cell size and density, and sometimes absent cell boundary markers. Additionally, sample quality and processing variations hinder obtaining reliable images, and the neglect of extracellular RNA complicates analysis. These issues make accurate, high-resolution analysis with current methods often unfeasible. Future updates of this comparative work will benefit from segmentation-free analysis^15,16^ to realize the full potential of emerging submicron iST technologies.

## Methods

### Patient material, ethics approval and consent for publication

The prostate cancer samples used in this study were obtained from patients undergoing radical prostatectomy, facilitated by the Australian Prostate Cancer BioResource. Patient consent was duly acquired in writing, adhering to the protocols sanctioned by the Human Research Ethics Committees of the University of Adelaide and St Andrew’s Hospital (Project approval numbers: H-2018-222 and #80). This consent encompassed the utilization of all de-identified patient data for publication purposes. Upon collection, the tumor tissues were promptly placed on ice, subsequently fixed in 10% neutral buffered formalin (NBF) for 24 hours, and then processed for paraffin embedding. For standard histological and pathological evaluations, the sections were stained with hematoxylin and eosin (H&E).

The specific patient tissue sample scrutinized in this study was biopsied from the left apex of the prostate. It has been categorized as infiltrating prostatic adenocarcinoma with a Gleason score of 3+4=7 (ISUP grade group 2). Notably, the sample lacks both Cribriform Gleason pattern 4 and intraductal carcinoma components. Additionally, there is no evidence of perineural invasion or invasion into the fat. Pertinent clinical notes for this patient include a PSA level of 6.6 µg/L and PIRDAS 4, affecting both the (L) apex and the (R) base (GG1 on surveillance).

### Tissue preparation and sequential sectioning

To ensure the highest quality of prostate tissue sections for our benchmarking study, we initially selected formalin-fixed, paraffin-embedded (FFPE) blocks containing biopsy material from four distinct patients who had recently undergone radical prostatectomy. All four blocks displayed exemplary morphology for benchmarking purposes, characterized by high cellularity and a balanced mix of tumor and normal tissue, as confirmed by H&E staining.

For each block, we face-cut and collected three consecutive 10 µm scrolls for the isolation and purification of total RNA using the Norgen Biotek FFPE RNA Purification Kit (Product ID 25300). We quantified the purified RNA with a NanoDrop One spectrophotometer (ThermoFisher Scientific). Subsequently, 5 ng of each RNA sample was analyzed using a 5200 Fragment Analyzer (Agilent) with the High Sensitivity RNA Assay kit (DNF-472). The DV-200 values, which indicate RNA integrity, varied across samples, ranging from 61.53% to 71.33%. We selected the block with the highest DV-200 value for all subsequent experiments in this study. The dimensions of the tissue in the selected block were approximately 7.5 mm by 5 mm.

To facilitate a comprehensive comparison of data from Xenium, CosMX, and snPATHO-seq platforms, we sectioned the chosen block as illustrated in Figure 1A. Specifically, we collected two 25 µm scrolls for snPATHO-seq and five 5 µm serial sections for Xenium and CosMX analysis (comprising two consecutive sections for each instrument and one additional section for H&E staining). Additionally, we prepared four more 25 µm scrolls for snPATHO-seq. Therefore, the sequential sections designated for each instrument were interspersed with a single 5 µm section for H&E staining, and these sections were flanked by scrolls for snPATHO-seq.

All sectioning procedures were conducted on a Leica RM2235 microtome, utilizing a fresh blade and ensuring thorough decontamination with RNAse AWAY (ThermoFisher Scientific). We have detailed the sectioning and subsequent slide preparation methods for Xenium and CosMx platforms below. The scrolls designated for snPATHO-seq and the single section reserved for H&E staining were stored at 4°C with desiccant until processed.

### Xenium slide preparation

Sections for Xenium were prepared according to the “Xenium In Situ for FFPE-Tissue Preparation Guide” (CG000578 Rev C, 10X Genomics) protocol. Briefly, 5 µm sections were cut, floated in an RNAse-free water bath, and carefully placed onto the Sample Area of a Xenium slide (PN-1000465) using a paintbrush. The slides were then placed in a drying rack at room temperature for 30 minutes to remove excess water, followed by a 3 hour incubation at 42°C on a Xenium Thermocycler Adapter plate, which was positioned atop the 96-well block of a C1000 Touch Thermocycler (BioRad) with the lid open. Subsequently, the slides were stored overnight at room temperature with a desiccant to allow further drying.

The next day, the Xenium slides were processed following the “Xenium In Situ for FFPE-Deparaffinization and Decrosslinking” protocol (CG000580 Rev C, 10X Genomics). In brief, the slides were placed again on a Xenium Thermocycler Adapter plate atop the 96-well block of a C1000 Touch Thermocycler (BioRad) with the lid open, and incubated at 60°C for 2 hours. After cooling, the slides underwent a series of immersions in xylene, ethanol, and nuclease-free water to deparaffinize and rehydrate the tissue.

Subsequently, the Xenium slides were assembled into Xenium cassettes (PN-1000566, 10X Genomics), which allow for the incubation of slides on the Xenium Thermocycler Adapter plate in a PCR machine with a closed lid for precise temperature control. The slides were processed using the “Xenium Slides and Sample Prep Reagents” kit (PN-1000460, 10X Genomics), starting with an incubation in a decrosslinking and permeabilization solution at 80°C for 30 minutes, followed by a wash with PBS-T.

The Xenium slides were then processed according to the “Xenium In Situ Gene Expression” user guide (CG000582 Rev D, 10X Genomics) for the remaining slide preparation steps. The slides were incubated at 50°C for 17 hours with the pre-designed gene expression probe set, “Xenium Human Multi-Tissue and Cancer Panel” (PN-1000626, 10X Genomics), which targets 377 individual human genes. This was followed by a series of wash and enzymatic steps, including a post-hybridization wash at 37°C for 30 minutes, a ligation at 37°C for 2 hours, and an amplification step at 30°C for 2 hours. After additional washing steps, the slides were treated with an autofluorescence quencher and a nuclei staining step.

### Xenium Analyzer setup and data acquisition

Processed Xenium slides, assembled in Xenium cassettes, were subsequently imaged using the Xenium Analyzer, following the guidelines provided in the “Xenium Analyzer User Guide: (CG000584 Rev B, 10X Genomics).” After completing a successful Instrument Readiness Test, run data, including Xenium slide ID numbers and pre-designed gene expression probe set information, was entered into the Analyzer. The instrument was equipped with the necessary consumables: Xenium Decoding Reagent Module A (PN-1000624, 10X Genomics), Xenium Decoding Reagent Module B (PN-1000625, 10X Genomics), Xenium Instrument Wash Buffer (PN-3001198, 10X Genomics), Xenium Sample Wash Buffer A (PN-30001199, 10X Genomics), Xenium Sample Wash Buffer B (PN-30001200, 10X Genomics), Xenium Probe Removal Buffer (PN-30001201, 10X Genomics), the Objective Wetting Consumable, the extraction tip, and a full rack of pipette tips.

The two Xenium slides/cassettes were then loaded into the instrument, initiating the Analyzer’s ‘sample scan’ process. This scan produced images of the fluorescent nuclei in each section. Upon completion, these images were utilized to determine the regions within the scan area to be included in the instrument’s comprehensive scan of the gene expression probe set. For both prostate sections, we essentially included the entire section in the full scan.

Post-run, the instrument was emptied of consumables and the Xenium slides carefully removed. Fresh PBS-T was added to each slide/cassette, and covered slides/cassettes were stored in the dark at 4°C until post-run H&E staining. Run data for each slide was copied from the Analyzer onto a Solid State Drive for further in-house analysis.

### CosMx slide preparation

Sections for CosMx were prepared and processed following the protocol “CosMX SMI Manual Slide Preparation” (MAN-10159-02, NanoString). Briefly, 5 µm sections were cut, floated in an RNase-free water bath, and then carefully placed onto SuperFrost Plus slides (Epredia J1800AMNZ) using a paintbrush. Since the slides imaged by the CosMx SMI are later assembled into a flow cell by applying a custom coverslip, it is crucial to place sections within a precisely defined area on each slide. A transparent template is used during sectioning to ensure correct placement.

The slides were then placed in a drying rack at 37°C for 2 hours to enhance section adherence to the glass. They were subsequently stored with a desiccant at 4°C until further processing. On the night before processing, the slides were baked in a drying rack at 60°C for 18 hours.

CosMx slides were then allowed to cool to room temperature and subjected to a series of xylene and ethanol immersions to deparaffinize the tissue. After drying at 60°C for 5 minutes, the slides were placed in a plastic Coplin jar containing 1X Target Retrieval Solution (CosMx FFPE Slide Prep Kit, Item # 121500006, NanoString), preheated to 100°C. The jar was then returned to 100°C for 15 minutes, followed by a quick rinse of the slides in room-temperature, nuclease-free water. The slides were washed in 100% ethanol and left to dry flat at room temperature for 30 minutes.

Incubation frames (CosMx FFPE Slide Prep Kit, Item # 121500006, NanoString) were placed on each CosMx slide, surrounding the section, to facilitate processing with small fluid volumes. Sections were permeabilized at 40°C for 30 minutes using a solution of pre-warmed proteinase K at a final concentration of 3 µg/ml in 1X PBS. The slides were then rinsed in 2X SSC, and fiducials at a final concentration of 0.001% (CosMx FFPE Slide Prep Kit, Item # 121500006, NanoString) were applied to the section and glass within the area covered by the incubation frame. After a wash with 1X PBS, each slide was post-fixed with 10% NBF, followed by a rinse in NBF stop buffer. The sections were then blocked with a solution of 100 mM NHS-acetate at room temperature for 15 minutes and washed with 2X SSC.

CosMx slides were prepared for probe hybridization using the CosMx Human Universal Cell Characterization Panel (Item # 121500002), which comprises a 950-target core panel of probes and a 50-target ‘add-on’ panel set. The core and add-on probes were denatured separately at 95°C for 2 minutes and immediately cooled on ice for 1 minute. Probes were then combined with Buffer R (CosMx FFPE Slide Prep Kit, Item # 121500006, NanoString) and applied to the slides for incubation at 37°C for 18 hours.

Post-hybridization, CosMx slides underwent two 25-minute washes at 37°C with a stringent wash buffer (1:1 formamide and 4X SSC), followed by two more washes in 2X SSC at room temperature. The slides were then treated with a blocking buffer and a nuclear stain (CosMx FFPE Slide Prep Kit, Item # 121500006, NanoString). After a rinse with 1X PBS, slides were incubated with antibodies from the CosMx Segmentation and Supplemental Markers Kit (Item # 121500020, 121500021, 121500022, NanoString), which included antibodies to CD298, B2M, PanCK, CD45, and CD68. Finally, the slides were washed three times with 1X PBS and then stored in 2X SSC until imaging.

### CosMx SMI setup and data acquisition

The setup and scan acquisition for the CosMx SMI instrument were conducted as outlined in the “CosMx SMI Instrument User Manual” (MAN-10161-02-1, Software Version 1.1, NanoString). We initiated a New Acquisition in the CosMx SMI Control Center web interface. Then, a CosMx RNA Imaging Tray (1,000 Plex 2 Slide Run) was taken out of storage at 4°C and allowed to reach room temperature for 30 minutes.

We carefully dried the CosMx slides in the areas outside the imaging region before placing each slide in the CosMx Flow Cell Assembly Tool. This was done by first lowering and then raising the tailgate. After removing the adhesive backing from a new flow cell coverslip, we carefully aligned the coverslip above the imaging area on the slide. The Assembly Tool was then firmly closed to attach the coverslip to the slide, thereby creating a functional CosMx flow cell. After removing the slide from the Assembly Tool, we gently applied 2X SSC into one of the two flow cell ports to hydrate the tissue section.

Next, we entered the flow cell configuration data into the Control Center interface. This included the barcode for each flow cell and the slide ID number. The prostate sections were scanned using Configuration C for the pre-bleaching profile and Configuration A for the cell segmentation profile.

We also entered details of the probe panel, cell segmentation, and supplemental marker into the flow cell configuration data for each section. After assembling the flow cells/slides, we loaded them into the instrument along with Buffer Bottles 1-4. Prior to placing Bottle 4 in the instrument, we added catalase and pyranose oxidase enzymes to it. The Control Center configuration was then verified, and following the addition of RNAse inhibitor to a single well of the CosMx reagent tray, the reagent tray was placed in the instrument as well. The instrument then performed a pre-run check and an initial scan of each tissue section.

Once the initial tissue scans were available in the Control Center, we selected a rectangular region around each tissue for a ‘pre-scan’. The pre-scan images, showing nuclear and cell segmentation marker staining, were used to select a contiguous region of 192 fields of view (FOVs) using the grid FOV placement tool. These FOVs completely encompassed the tissue areas on each slide. After approving the FOV selections for each slide, we initiated the 1,000-plex probe set scans for both slides.

CosMx scan data is automatically uploaded to NanoString’s cloud-based AtoMx^TM^ Spatial Analysis Platform during the run (see the CosMx Data Analysis manual (MAN-10162-02, Software Version 1.1, Nanostring) for more detail. Briefly, once the run was complete and scan data fully uploaded to AtoMx^TM^, a study was created for each CosMx scan. Study creation in AtoMx^TM^ allows the user to create and run a set of data analysis pipelines, including a data export pipeline. This workflow was used to export a variety of CosMx data, including TileDB arrays and Seurat objects for further in-house analysis.

### H&E staining of sections; imaging of H&E stained sections

For H&E staining of the virgin 5 µm prostate tissue section that lies between the Xenium serial sections and those of CosMx, the slide was baked at 60 °C for 60 minutes and then processed through the Dako Progressive H&E protocol (Agilent) and using Dako Harris Hematoxylin, Dako Bluing Buffer and Dako Modified Eosin Y. For post-run H&E staining of Xenium slides, the protocol from “Post-Xenium Analyzer H&E Staining” (CG000613 A, 10X Genomics) was followed exactly. In brief, Xenium slides were first incubated in 10 mM sodium hydrosulphite at room temperature for 10 minutes to remove the autofluorescence quencher from the sections, rinsed three times in Milli-Q water, and then processed through a relatively standard protocol utilizing Mayer’s hematoxylin, eosin Y and bluing solution. (precise steps and timing are found in the 10X Genomics document). After the last xylene wash, the virgin and post-Xenium sections were dried for 15 minutes and then coverslipped using 18 x 18 mm #1.5 cover glass and Dako toluene-free mounting media (CS705).

Post-run H&E staining of CosMx slides also utilized the precise steps and timing of the 10X Genomics H&E staining protocol, except for the initial sodium hydrosulphite treatment. As the post-run CosMx sections remained encased in their flow cells, we developed a silicone gasket seal that could be applied to one of the two inlet/outlet ports of the CosMx flow cell, and which could provide a tight seal between the flow cell and a P1000 pipette tip. This gasket allowed us to rapidly flow fluids over the CosMx sections, and thus we used each flow cell as a mini-chamber for the H&E staining reagents. At the end of H&E staining the last xylene wash was completely removed from each flow cell, and the sections were allowed to dry overnight. A tungsten carbide glass scribe was then used to cut the flow cell glass just inside of the adhesive boundary of the flow cells. The cut flow cell glass was carefully removed from the slides and the sections coverslipped as above but with a smaller 9 x 18 mm cover glass.

H&E stained sections were scanned in semi-automatic mode on a NanoZoomer 2.0-HT, using a 40X objective with manual focus point selection. Xenium H&E images were also imaged on a Zeiss AxioObserver 7 using a 20X objective. Zeiss .zvi images were then exported as pyramidal ome.tif files using QuPath-0.4.0 ^19^, and then imported into Xenium Explorer.

### Single-nucleus RNA-seq on FFPE tissue (snPATHO-seq)

Two tissue sections, each 25-30 μm in thickness, were initially washed three times with 1 mL of xylene for 10 minutes to remove paraffin. They were then rehydrated through a series of 1 mL ethanol washes, each lasting 1 minute: two consecutive washes in 100% ethanol, followed by washes in 70%, 50%, and then 30% ethanol. The initial xylene wash was performed at 55°C. The sections were subsequently briefly rinsed with 1 mL of RPMI1640 (Gibco).

Tissue disruption began with physical homogenization using a pestle in 100 μL of a digestion mixture. This mixture consisted of 1 mg/mL Liberase TM (5401119001, Roche), 1 mg/mL Collagenase D (11088858001, Roche), and 1 U/μL of RNase inhibitor (RiboLock RNase Inhibitor, EO0382, Thermo Fisher Scientific) in RPMI1640. The mixture was then brought up to a 1 mL volume and incubated at 37°C for 45-60 minutes for digestion.

For nuclear extraction, the pre-digested tissue was treated with 1x Nuclei EZ lysis buffer (NUC-101, Sigma-Aldrich) containing 2% BSA. The disintegration process was further facilitated by pipetting with a P1000 pipette. The liberated nuclei were filtered through a 70 μm mesh, washed twice with 1x PBS supplemented with 1% BSA, and once with a 0.5x PBS solution containing 0.02% BSA. They were then resuspended in the latter solution and re-filtered through a 40 μm mesh (pluriStrainer Mini 40μm, 43-10040-50, pluriSelect). The final nuclear count was determined using a LUNA-FX7 cell counter (AO/PI viability kit, F23011, Logos).

Gene expression libraries were prepared using the Chromium Fixed RNA Kit, Human Transcriptome (1000474 or 1000475, 10x Genomics), following the user guide (Chromium Fixed RNA Profiling, CG000477 - RevB, 10X Genomics). We performed additional optimizations in the pre-amplification and indexing cycles, specifically tailored for nuclei derived from FFPE samples. The sample was divided into four identical aliquots, each containing over 400,000 nuclei, and subjected to a 20 hour hybridization with the BC01-BC04 probe set. Subsequent to three pooled washes with the Post-Hyb Buffer as recommended in the guide, an extra wash step was included. After the post-hybridization washes, the nuclei were resuspended in the Post-Hyb Resuspension Buffer, recounted using AO/PI staining on the Luna FX-7 instrument (Logos Biosystems), and loaded onto two lanes of a Chip Q/Chromium X for capture (targeting 40,000 cells/lane), in accordance with the guide’s procedures. A 9-cycle pre-amplification was performed, followed by indexing PCR cycles as recommended for PBMC and nuclei in the guide, but with an additional two cycles.

### Processing snPATHO-seq data

The transcript count matrices from cellranger were imported and processed with Seurat v5.0.1^20^. Prior to quality control filtering, doublets were inferred in each library using scDblFinder v1.14.0 with a doublet rate of 10%, approximating the expected rate from the multiplexed fixed RNA profiling. Examining the distribution of UMI and features counts per nuclei, there was a prominent spike of droplets with low counts. We first confirmed this population lacked distinct expression patterns and likely reflected debris or ambient RNA. Nuclei annotated as doublets and those with fewer than 1000 detected genes were filtered, and we confirmed that no specific cell types were lost compared to unfiltered data. Normalization and feature selection was performed with SCTransform with default parameters prior to dimensionality reduction with principal component analysis. The first 30 components were then used as input to produce a UMAP embedding (RunUMAP) and for Louvain clustering.

To annotate the data, scRNA-seq data of human prostate from Tuong *et al*.^10^ was used as a reference to transfer cell type labels onto the snPATHO-seq data. This was performed using Seurat’s label transfer method, identifying anchors across datasets based on the first 30 components of the reference’s PCA embedding. The purity of annotations within individual clusters was high, however the reference data did not include annotations for several prominent mesenchymal cell types. For example, smooth muscle, skeletal muscle, fibroblasts, and pericytes were all annotated as “Fibroblast”. We supplemented the annotations with these cell types based on their marker expression (Fibroblast, *PDGFRA*; Pericyte, *PDGFRB/ACTA2*; Smooth muscle, *MYH11*; Skeletal muscle, *TTN*). The final annotations reflect the prominent annotation of each cluster following label transfer.

### Processing of Xenium and CosMx data

To perform a fair comparison of data from both platforms, only cells with no detected transcripts were removed from the data following the Xenium Onboard Analysis or the AtoMx^TM^ processing workflow. Effectively all analysis, including cell type annotation, was performed on raw transcript counts. UMAP embeddings were generated for the purpose of visualization alone, first normalizing the data with SCTransform, performing principal component analysis, and then using the first 30 components to produce the UMAP embedding. Visualizations of transcript localization and segmentation boundaries were generated using Seurat’s ImageDimPlot() function.

### Automated annotation of spatial transcriptomics data

Cells from both Xenium and CosMx were annotated with InSituType v1.0.0^21^, using the snPATHO-seq data as a reference. We produced reference transcript counts for each cell type by taking the average UMI counts from the corresponding cell type in the snPATHO-seq data. The average counts for the negative control probes for each assay were also calculated. These profiles were then used as input for the insitutypeML function along with raw counts from the spatial transcriptomics data for annotation. For some analyses, annotations for the multiple epithelial and muscle cell types were simplified as either “Epithelial” or “Muscle”.

We note that we also explored Seurat’s label transfer method. While annotations were fairly consistent for the Xenium data, the annotations of the CosMx samples were heavily skewed towards epithelial annotations. InSituType annotations better reflected the tissue structure and were used for all analyses.

### Spatial autocorrelation

Moran’s I was used to evaluate the spatial patterning of transcripts throughout the tissue. To compute this, we used the method implemented in the R package Voyager v1.2.7^22^. Briefly, transcript count and spatial coordinate for each cell were used to construct a SpatialFeatureExperiment. After normalizing the counts with logNormCounts, findSpatialNeighbors was used to construct a nearest neighbor graph (k=20) based on the inverse distance weight (dist_type = “idw”). This graph was then used to calculate global Moran’s I values for each gene using the runMoransI function.

## Authors Contributions

L.G.M. and J.T.P. designed the project, interpreted the results, and wrote the manuscript. D.P.C. conducted all the bioinformatic analyses, interpreted the results, made all the figures, and wrote the manuscript. K.B.J. and K.W. conducted the assays and assisted in writing the manuscript. N.K.R. and L.M.B. supplied the samples. M.J.R., F.S.-D., M.Z., I.S.V., S.R.V.K., J.L.W., L.M.B., and N.B. critically reviewed and provided feedback on the manuscript.

## Acknowledgements

We extend our gratitude to the staff at the University of Adelaide Histology facility for their expertise in tissue sectioning. We also thank the Australian Genome Research Facility team for their rapid sequencing turnaround. Our appreciation goes to the South Australian Genomics Centre for their support during this study. Additionally, we would like to acknowledge the Field Application Specialists from 10X Genomics and NanoString Technologies for their exceptional support. We thank Prof. Alistair Forrest for his critical review of the manuscript. Special thanks to 10X Genomics and NanoString Technologies Support Teams for their training and guidance in overcoming some of the technological challenges encountered during the project.

## Disclaimers

L.G.M., J.T.P., D.P.C., D.P.K., K.B.J., K.W., M.J.R., F.S-D., N.K.R., M.Z., I.S.V., S.R.V.K., L.M.B., J.L.W. and N.B. declare no professional or financial affiliations with 10X Genomics or NanoString Technologies. During the conduct of this study, L.G.M. served as a Scientific Advisor for Millenium Sciences (no longer in this role), Omniscope, and ArgenTag. N.B. is Chief Scientist at Deepcell. S.R.V.K. is the founder of and a consultant for Faeth Therapeutics and Transomic Technologies. None of the authors received any form of payment or compensation from these companies. Consumables used in this study from both companies were purchased at full price, acquired at a discounted rate, or provided free of charge, although not specifically for this study. Subscri ption to AtoMx^TM^ was provided at no cost by NanoString Technologies.

## Data Availability

Data will be made publicly available shortly and directions for access will be added to the GitHub repository listed below. If access is needed prior to this, please contact the authors.

## Code Availability

Code to reproduce this analysis is available at https://www.github.com/dpcook/spatial_benchmark

